# Temporal Lobe Activation Predicts Episodic Memory Following Traumatic Brain Injury

**DOI:** 10.1101/2020.12.01.407064

**Authors:** Abbie S. Taing, Matthew E. Mundy, Jennie L. Ponsford, Gershon Spitz

## Abstract

The temporal lobes are critical for episodic memories and are preferentially affected following a traumatic brain injury (TBI). As such, episodic memory difficulties are common following TBI; however, the underlying neural changes that precipitate or maintain these difficulties in the early phase of recovery remains poorly understood. Here, we use functional magnetic resonance imaging (fMRI) to interrogate the relationship of temporal lobe activation in response to face, scene, and animal stimuli. Twenty-five patients with moderate to severe TBI were recruited an average of 2 months’ post-injury and compared with 21 demographically similar healthy controls. Findings indicate that memory for faces was preferentially impaired, compared to scene and animal stimuli. Decreased activity in temporal lobe structures was present for both face (right transverse temporal gyrus) and scene stimuli (right fusiform gyrus), but not for animals. Greater activation in these structures was associated with better long-term recognition. These findings provide evidence to suggest that TBI: a) preferentially affects memory for complex stimuli such as faces and scenes, and b) causes aberrant neuronal processes despite lack of evidence of significant impairment in behavioural performance. The mechanisms underpinning these findings are discussed in terms of differences in strategy use and reduced neural efficiency.

## Introduction

Traumatic brain injury (TBI) can cause focal and diffuse disruption to multiple brain systems. Pathology is most widely observed in frontal and temporal cortices (Bigler 2001), and as such impairments in processing speed, attention, executive function, and memory are most common (Draper and Ponsford 2008; Azouvi et al. 2017). Invariably, all individuals with a moderate or severe TBI will experience an initial transient period of impaired consciousness with amnesia and general confusion known as post-traumatic amnesia (PTA; Tittle and Burgess 2011). Although most individuals emerge from PTA, many report ongoing difficulty with so-called episodic memories (Vakil 2005; Rabinowitz and Levin 2014). Episodic memories involve the ability to learn, store, and retrieve information about personal experiences (Tulving 2002; Moscovitch et al. 2016). Impairment of episodic memory can interfere with crucial skills such as new learning and task completion, and therefore can significantly limit functional independence and productivity (Nakase-Richardson et al. 2011).

Neuroanatomically, episodic memory is supported by a brain network spanning several regions in the frontal and parietal cortices (Wagner et al. 2005; Eichenbaum 2017). However, structures contained within the temporal lobes are arguably the most crucial for particular aspects of episodic memory (Dickerson and Eichenbaum 2010; Moscovitch *et al.* 2016). The temporal lobes house the hippocampus, and the entorhinal, perirhinal, and parahippocampal cortices – collectively known as the medial temporal lobes (Simons and Spiers 2003; Graham et al. 2010). The hippocampus and other medial temporal structures are involved in the encoding of episodic stimuli, and modulation of activity in these regions has been found to predict subsequent recollection of the encoded stimuli (Cameron et al. 2001).

The temporal lobes are preferentially affected after TBI (Bigler 2001). The location of the temporal lobes and their proximity to the middle cranial fossa makes them highly susceptible to acceleration-deceleration forces present during injury (Barlow 2013; Daneshvar and McKee 2015). Structures within the temporal lobes, such as the hippocampus and other limbic structures, are particularly susceptible to hypoxic insults and excitotoxicity effects (Dickerson and Eichenbaum 2010). As such, atrophy in the hippocampus and fornix is common following TBI (Bigler et al. 1996), and has been associated with impaired memory (Ariza et al. 2006).

Functional imaging studies in the TBI population have also implicated the temporal lobes in impaired episodic memory. Individuals with TBI tend to display increased activity in various regions, including the temporal lobes, during encoding of episodic stimuli when compared with healthy controls (Russell et al. 2011; Arenth et al. 2012; Gillis and Hampstead 2015). One limitation of these previous studies is that most have failed to use paradigms that may reflect the type of episodic memory deficits that individuals with TBI may experience in their daily life. Both Russell *et al.* (2011) and Arenth *et al.* (2012) used a mixture of line drawings of pictures and shapes, words, and letter strings, and only short-term retention was assessed. Gillis and Hampstead (2015) addressed these methodological shortcomings by using realistic images of common objects and assessed long-term retention outside of the scanner. However, participants recruited into their study were in the chronic stages of recovery (range 1 to 13 years). Characterisation of memory for ecologically relevant episodic stimuli following TBI is lacking for the early period of recovery.

Here, we rectify this gap by investigating the extent to which memory for common stimuli is impaired following TBI, focussing specifically on the temporal lobe. Processing of common stimuli such as faces, scenes, and animals has been found to recruit specialised temporal lobe structures (Downing et al. 2006; Mundy et al. 2009). Exposure to faces robustly activates regions in the middle fusiform gyrus (‘fusiform face area’), the lateral inferior occipital gyrus (‘occipital face area’), and right superior temporal sulcus (Hoffman and Haxby 2000; Kesler et al. 2001; Haxby et al. 2002); exposure to scenes activates the posterior parahippocampus (‘parahippocampal place area’; Epstein and Ward 2010); and exposure to animals recruits activity in the bilateral fusiform gyrus (Rogers et al. 2005; Downing *et al.* 2006). Although some degree of impairment is expected given the high prevalence of temporal pathology and specialised processing of various stimuli in this area, it is possible that not all stimuli are equally impaired following injury. Past studies of amnestic patients have demonstrated stimulus-sensitive impairments for complex stimuli such as faces and scenes (Taylor et al. 2007; Mundy et al. 2013). In the TBI population, impairment of face recognition has also been previously documented (Valentine et al. 2006).

The temporal lobes play a critical role in episodic memory, which is frequently impaired following TBI. Therefore, the present study specifically focusses on the temporal lobes to examine how episodic memory impairments may affect aspects of everyday memory in the early phase of recovery. We used an fMRI task to measure temporal lobe processing during encoding of three stimuli of varying complexity: faces, scenes, and animals. Recognition memory was subsequently probed in an out-of-scanner behavioural task. In line with previous studies (Russell *et al.* 2011; Arenth *et al.* 2012; Gillis and Hampstead 2015), we hypothesised that the TBI group would display greater temporal lobe activation during stimulus encoding compared to healthy controls. Furthermore, we hypothesised that greater activity during encoding would be associated with poorer recognition task performance. Lastly, we hypothesised that individuals with TBI would be most impaired for complex stimuli such as faces and scenes.

## Materials and Methods

### Participants

Twenty-five patients (17 males, 8 females) who had sustained moderate to severe TBI, determined prospectively using the Westmead Post Traumatic Amnesia Scale (Shores et al. 1986), were recruited from the Acquired Brain Injury Ward at Epworth Hospital (Richmond, Victoria) after emerging from post-traumatic amnesia (*M* = 2.16 months, *SD* = 1.48 months, range = 0.69 – 6.64 months; Table 1 and Supplementary Table 1). TBI patients predominantly had prefrontal and temporal pathology (Fig. 1). Exclusion criteria included age < 18 or > 75 years, prior history of TBI or other neurological conditions, significant psychiatric or substance abuse history, and MRI contraindication. Twenty-one healthy controls (13 males, 8 females) of similar age, sex, and education were also recruited (Table 1). There were no significant group differences on any of the demographic variables (*P* > 0.05). Due to a technical error during data collection, a subset of the sample (6 TBIs and 8 healthy controls) was excluded from the MRI analyses due to poor coverage of the temporal lobes. Additionally, 1 healthy control participant was excluded due to excessive movement in the scanner. Written informed consent was provided by all participants in accordance with the Declaration of Helsinki. This study was approved by Monash Health Human Research Ethics Committee.

**Table 1.** Demographic information and clinical characteristics of the TBI and healthy groups.

| Demographic variables | Traumatic brain injury | Healthy controls, |
| --- | --- | --- |
|  | Mean (SD) | Mean (SD) |
| Age (years) | 38.36 (16.82) | 37.29 (17.89) |
| Sex (male/female) | 17/8 | 13/8 |
| Education (years) | 13.82 (3.27) | 14.19 (2.57) |
| Time since injury (months) | 2.16 (1.51) | - |
| PTA (days) | 22.46 (14.63) | - |
| GCS (lowest) | 9 (4.37) | - |

GSC = Glasgow Coma Scale; PTA = post-traumatic amnesia. Note: PTA duration were available for $n = 24$ patients.

**Figure 1.**
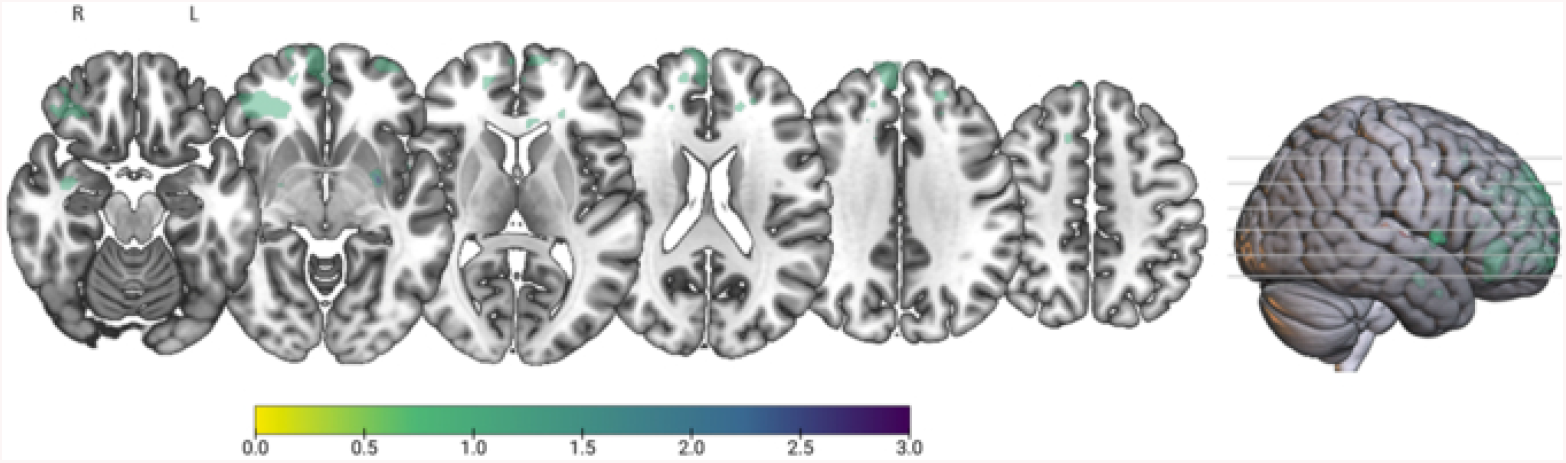
Lesion overlay plot of all TBI participants. Maps were overlaid on a T1 template in MNI space.

### Episodic Memory Paradigm

The episodic memory paradigm is a task adapted from Mundy et al. (2012; 2013) to probe episodic memory encoding and recognition (Figure 2). Participants were presented images of faces, scenes, and animals while in the scanner. They were instructed to respond, using trigger buttons, to each stimulus based on set criteria to ensure attention was maintained throughout the task (e.g. decide whether an animal is shorter or taller than a human man; whether the face is male or female; whether a scene looks hot or cold). The fMRI task consisted of six blocks, each block comprising 20 images from the three stimulus categories. This task comprised 60 unique stimuli, each presented twice throughout the session. Each stimulus was presented for 3 seconds, followed by a 3 second inter-stimulus interval. Five rest blocks were presented after the 1^st^ – 5^th^ experimental blocks.

**Figure 2.**
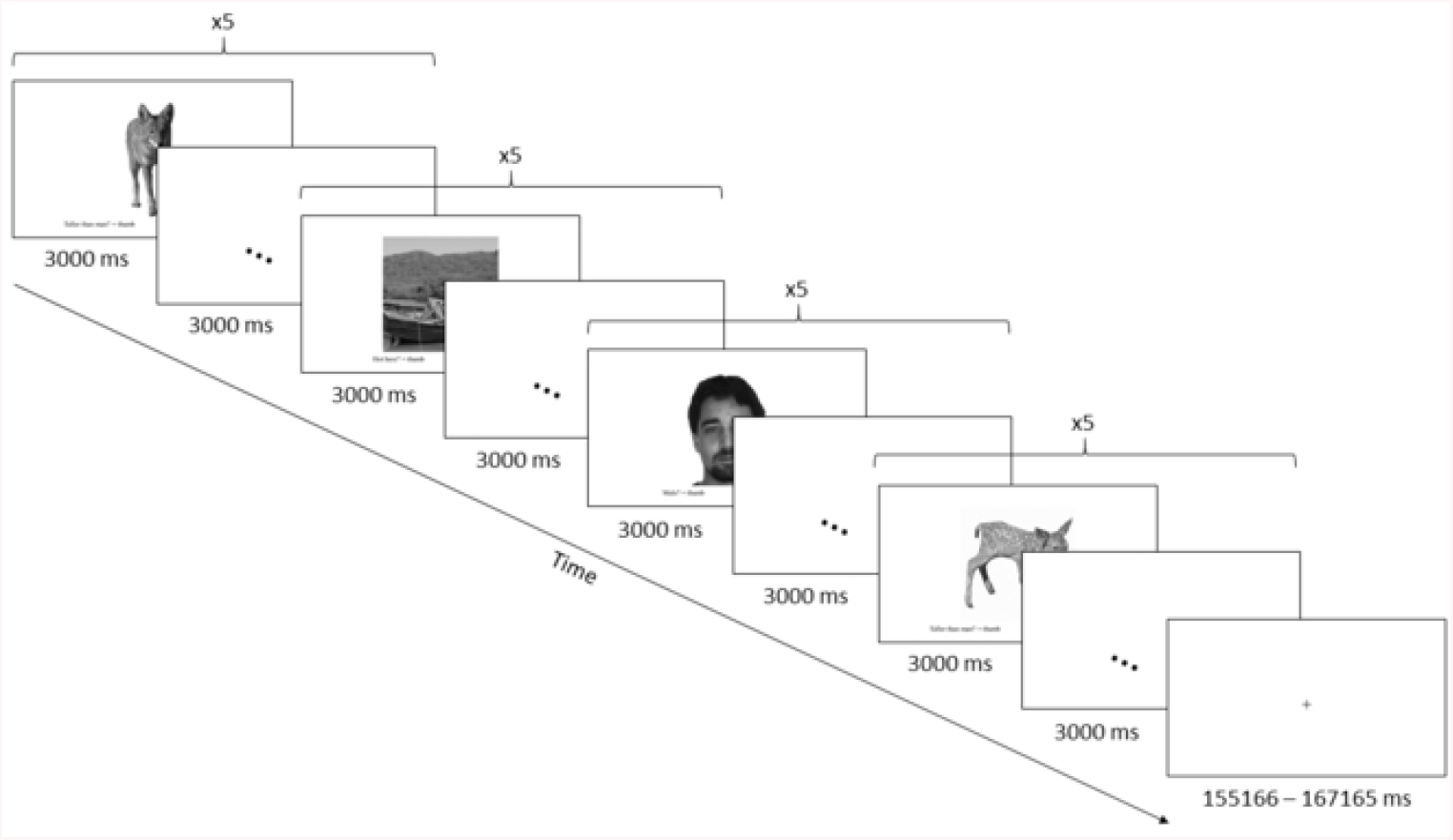
Schematic diagram of the episodic encoding task. Participants were presented 5 images from a stimulus category (i.e. faces, scenes, and animals) and were instructed to response to the various stimuli on screen based on set criteria. Each stimulus was presented for 3 s, followed by an inter-stimulus duration of 3s.

Memory of the stimuli presented in the fMRI task was probed in an out of scanner task. Recognition of the episodic stimuli was assessed by instructing participants to classify stimuli as “old” (i.e. images seen during the fMRI task) or “new” (i.e. images that were not presented during the fMRI task). Participants rated a total of 60 old and 60 new stimuli, each presented twice (240 stimuli in total). To assess retrieval ‘confidence’, images were rated on a scale from “definitely old” to “definitely new”. Responses were considered correct/incorrect regardless of confidence level (secondary analysis indicated no significant differences in confidence rating between groups – see “Confidence Rating Analysis” in Supplementary for further detail). Performance for this task was assessed using accuracy and reaction times.

### MRI Acquisition

Structural and functional MR images were acquired in two clinical scanners, using 3.0 Tesla Siemens Magnetom Skyra scanners and a 20-channel head coils. Functional images were acquired using single-shot gradient-echo planar imaging (EPI) with the following parameters: repetition time (TR) = 2.75 s; echo time (TE) = 30 ms; flip angle = 90°; 220 × 220 matrix; voxel size = 3.4 × 3.4 × 3.0 mm. A high-resolution 3D T1-weighted image covering the entire brain was also acquired for anatomical reference (TR = 2.3 s; TE = 2.32 ms; flip angle = 8°; 236 × 350 matrix; voxel size = 0.9 × 0.9 × 0.9 mm). Due to reduced brain coverage (due to using clinical scanners), and a-priori hypotheses, we focused on the temporal lobes.

### Statistical analysis

#### Behavioural and Demographic Data

Behavioural and demographic data were analysed using R version 3.6.0 (R Core Team, 2019). Two-tailed independent samples t-tests were used to examine group differences on the demographic variables including age, sex, and years of education. Behavioural data were screened for normality, transformed (if necessary), and assessed for violation of statistical assumptions prior to analysis. Outcome measures were analysed using linear mixed models to account for clustering or non-independence of measures within participants. Memory performance was assessed using dprime and reaction time. Dprime measured task accuracy, accounting for the signal (hits) and noise (false alarms). Reaction times were generated by obtaining the average reaction time per stimulus category. Task accuracy was assessed by modelling stimulus category, group, and their interaction (stimulus category x group) as fixed effects, and participant as a random effect. For reaction time, the data were inversely transformed, and a model was fitted with stimulus category, group, and an interaction (stimulus category x group) as fixed effects, and participant as a random effect. Where appropriate, post-hoc analyses were conducted using two-tailed t-tests with Bonferroni correction.

### Imaging Data

#### MRI Preprocessing

Prior to preprocessing, lesions were manually segmented using MRIcron (http://www.mricro.com/mricron). Preprocessing was performed using fMRIPrep 20.0.0 (Esteban et al., 2018) and involved the application of the following step: undistortion of EPI data, realignment, normalisation, and estimation of confounds. Further information about the MRI preprocessing can be accessed in the Supplementary (see “Detailed MRI Preprocessing”).

#### fMRI Analysis

fMRI data were analysed using FSL’s FEAT version 6.0.2 (FMRIB’s Software Library, www.fmrib.ox.ac.uk/fsl). In the first level analysis, contrasts between each stimulus category (i.e. animals, faces, scenes) and the rest blocks were generated for each participant. The onset times for each contrast corresponded to the first stimulus presentation of each category and were 27 seconds in duration. To reduce motion-related artifacts, additional regressors using a modified method of the anatomical CompCor that explained 50% of the variance were also included in the first level model (Muschelli et al. 2014). Differences in brain activation of categories of episodic stimuli were assessed in a 2 (group: TBI vs. healthy controls) x 3 (stimulus category: faces, scenes, and animals) factorial design using FLAME 1 +2 mixed effects with automatic outlier de-weighting. An additional explanatory variable was also added at the group level to control for the acquisition of images from two scanners. Given our hypothesis regarding structures in the temporal lobes and their role in learning and processing of category-specific stimuli, an a-priori region of interest (ROI) mask of the temporal lobe was generated using the MNI Structural Atlas (Supplementary Figure 1) and used in the group level analysis. Imaging findings are reported using a cluster level threshold of *Z* > 3.1 and a family wise error cluster correction threshold of *P* < 0.05. FEAT contrast of parameter estimates (COPE) were extracted from significant clusters at the group level. To investigate differences in BOLD response, two-tailed independent samples t-tests were conducted using COPE values. Finally, association between BOLD response and behavioural performance on the episodic recognition task were assessed using Pearson correlations.

### Data Availability

All data supporting the findings of this study can be requested from the corresponding author.

## Results

### Episodic Memory is Selectively Impaired for Face Stimuli

First, we investigated whether recognition of episodic memories was impaired following TBI. To do this, we performed a linear mixed model assessing memory recall accuracy (old vs. new, as measured using dprime) on the episodic retrieval task. Overall, the TBI group demonstrated significantly poorer recognition accuracy than healthy controls (95% CI, 0.06 – 0.74; *P* = 0.023). Post-hoc analyses indicated that the group difference was driven by a significant difference in accuracy for faces (95% CI, 1.10 – 1.50; *P* = 0.028). There was a trend for lower accuracy for scenes, however, this did not reach statistical significance (95% CI, 0.93 – 1.20; *P* = 0.140). There was no significant difference between the groups in accuracy for animals (95% CI, 1.11 – 1.27; *P* = 0.383).

### Reaction Times were Quickest for Face Stimuli

Next, we investigated whether reaction time was significantly different between the groups and whether it varied according to the type of stimulus (Figure 3). Overall, reaction time was greater for individuals with TBI compared to healthy controls, irrespective of stimulus type (95% CI, 0.08 – 0.25; *P* < 0.001). Post-hoc analyses revealed that the TBI group was significantly slower than healthy controls in responding to faces (95% CI, 0.48 – 0.64; *P* < 0.001), scenes (95% CI, 0.43 – 0.58; *P* = 0.001), and animals (95% CI, 0.42 – 0.58; *P* = 0.001). Across both groups, reaction time also varied depending on the stimulus category: the TBI group was quicker to respond to faces than scenes (95% CI, 0.43 – 0.48; *P* < 0.001) and animals (95% CI, 0.42 – 0.48; *P* < 0.001); similarly, healthy controls were quicker to respond to faces than scenes (95% CI, 0.58 – 0.64; *P* = 0.003) and animals (95% CI, 0.58 – 0.64; *P* = 0.001). Given this pattern of results, we further investigated whether the poorer performance for faces in the TBI group was driven by a speed-accuracy trade-off. To do this, we included reaction time as a covariate in the linear mixed model. Results indicated that participants with TBI still performed significantly poorer than healthy controls after controlling for reaction time (95% CI, 1.09 – 1.55; *P* = 0.028), suggesting that this pattern of performance was not due solely to a speed-accuracy trade-off.

**Figure 3.**
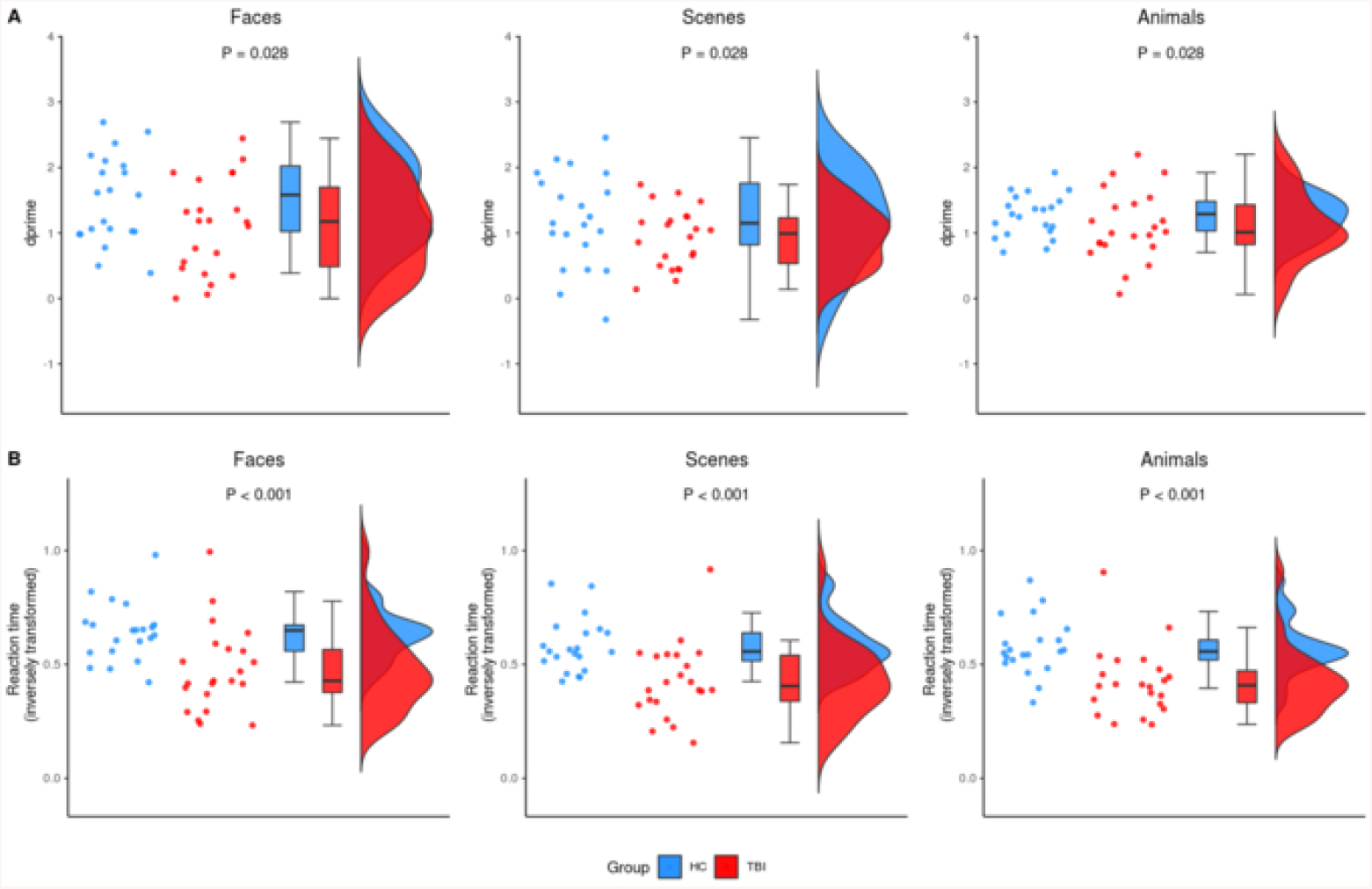
Behavioural results for the episodic retrieval task. A) Plots representing accuracy according to stimulus category, as measured using dprime (higher values denote better performance). The TBI group had significantly poorer accuracy than healthy controls when retrieving faces (*P* = 0.028). There was a trend for lower accuracy for scenes, although this did not reach statistical significance (*P* = 0.140). There was no significant difference in accuracy for animal stimuli (*P* = 0.383). B) Plots representing reaction time according to stimulus category (note: reaction time was inversely transformed; higher values denote faster performance). As expected, the TBI group was slower than healthy controls when responding to the various stimuli (*P* < 0.05).

**Figure 4.**
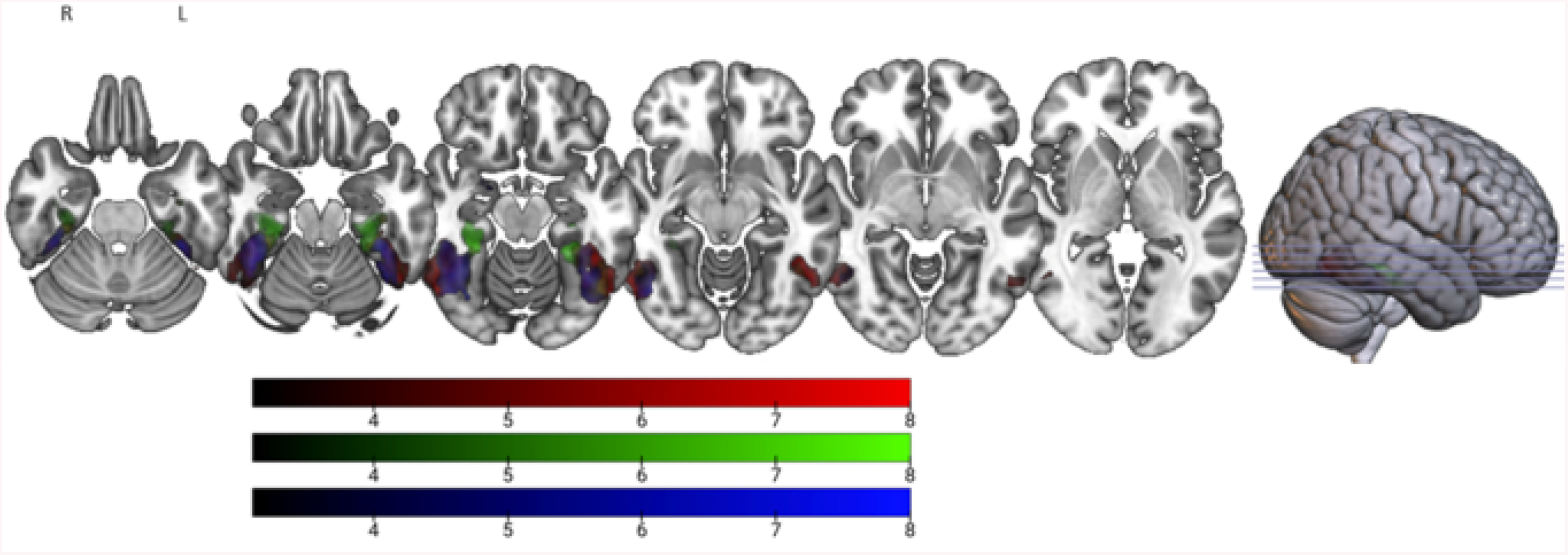
Overall functional activity elicited during the episodic encoding task for the whole sample. Significant clusters during encoding of faces (blue), scenes (green), and animals (red).

### fMRI Task Activates the Stereotypical Regions Underpinning Encoding of Episodic Stimuli

To demonstrate that our task elicited activations in stereotypical areas involved with the canonical network that support encoding of episodic stimuli, we first included all participants in an analysis looking at the average activation for each stimulus category (i.e. faces, scenes, and animals). During encoding of face stimuli, significant clusters were noted in face-selective areas including the right inferior occipital gyrus and left/right fusiform gyrus (Hoffman and Haxby 2000; Kesler *et al.* 2001; Haxby *et al.* 2002), as well as the right hippocampus. During encoding of scene stimuli, significant clusters were noted in scene-selective area of right parahippocampal gyrus (Epstein and Ward 2010), as well as the left fusiform gyrus. Finally, during encoding of animal stimuli, significant clusters were noted in animal-selective area of the right fusiform gyrus (Rogers *et al.* 2005; Downing *et al.* 2006), as well as the left inferior occipital gyrus.

### TBI Patients Show Reduced Right Transverse Temporal Gyrus Activation During Face Processing

Consistent with the behavioural results, group differences on imaging were apparent during encoding of faces. TBI participants showed reduced activation in the right transverse temporal gyrus extending to the planum temporale compared to healthy controls (Figure 5).

**Figure 5.**
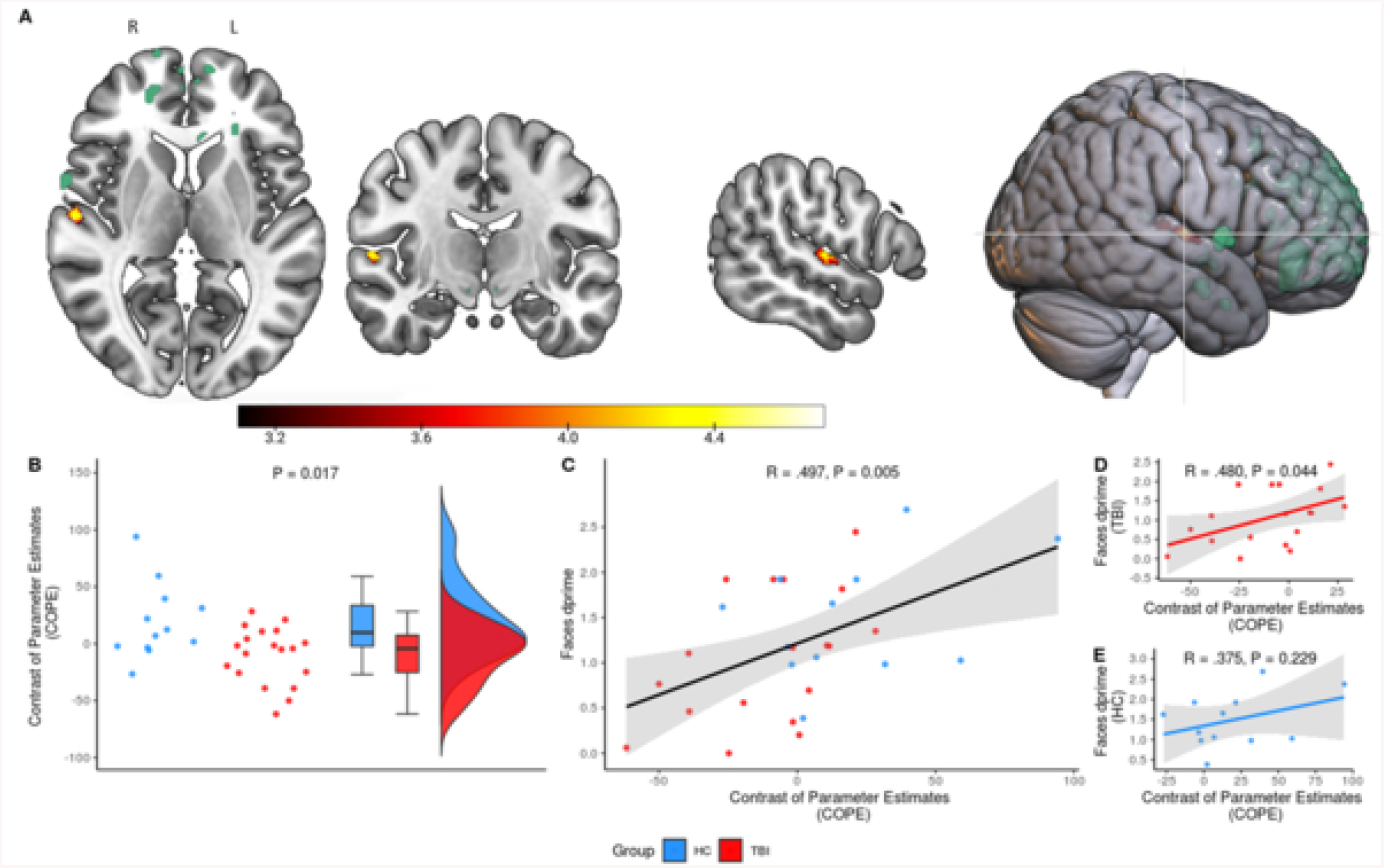
Reduced functional activity during encoding of faces associated with behavioural performance. A) The TBI group demonstrated a reduced response in the right transverse temporal gyrus extending to the planum temporale during encoding of faces compared to healthy controls. There was no overlap between this cluster and lesions overlap (green). Note: cluster was dilated to increase visibility. B) Plot of the COPE values extracted from the significant cluster in the right transverse temporal gyrus extending to the planum temporale. The difference in COPE values between the TBI group (red) and healthy controls (blue) was significant (*P* = 0.017). C) Overall, there was a significant association between COPE values and the dprime scores for face stimuli (*P* = 0.005). However, further investigation indicated that a significant association was only apparent for D) TBI group (*P* = 0.044), and not E) healthy controls (*P* = 0.229).

FEAT analysis COPE values were extracted for the significant cluster. We first investigated whether there was a significant difference in BOLD response between groups using an independent samples t-test. As expected, TBI patients displayed lower COPE values (*M* = −9.90, *SD* = 24.79) compared to healthy controls (*M* = 19.07, *SD* = 33.04), *t*(18) = −2.61, P < 0.017. We further examined whether there was an association with behavioural performance on the episodic recognition task using Pearson correlations. Overall, there was a moderate positive relationship between COPE values and the dprime scores for face stimuli, *r*(28) = 0.497, *P* = 0.005. Follow-up correlations indicated that a significant correlation was only apparent for the TBI group, *r*(16) = 0.480, *P* = 0.044, and not healthy controls, *r*(10) = 0.375, *P* = 0.229.

### TBI Patients Display Reduced Right Fusiform Gyrus Activity During Scene Processing

Despite a non-significant difference between groups for scene recognition, we found reduced activation in the right posterior fusiform gyrus for the TBI group in comparison to healthy controls during scene encoding (Figure 6).

**Figure 6.**
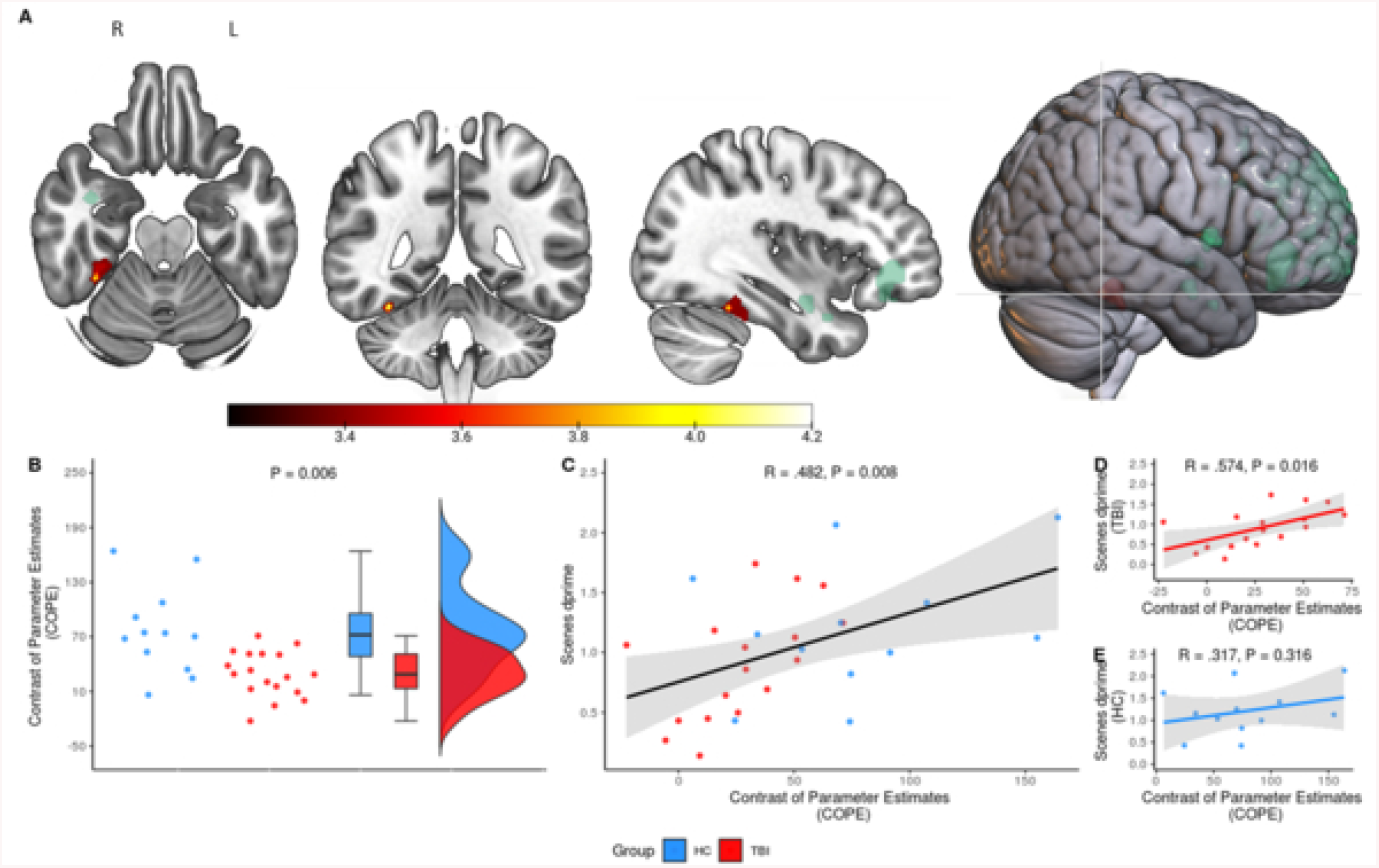
Reduced functional activity during encoding of scenes was associated with behavioural performance. A) In comparison to healthy controls, the TBI group demonstrated reduced activity in the right posterior fusiform gyrus encoding of scene stimuli. There was no overlap between this cluster and lesions overlap (green). Note: cluster was dilated to increase visibility. B) Plot of the COPE extracted from the significant cluster in the right posterior fusiform gyrus. The difference in COPE between the TBI group (red) and healthy controls (blue) was significant (*P* = 0.006). C) Overall, there was a signification association between the COPE and the dprime for scene stimuli (*P* = 0.008). However, further investigation indicated that a significant association was only apparent for D) TBI group (*P* = 0.044), and not E) healthy controls (*P* = 0.229).

TBI patients showed lower COPE values (*M* = 29.26, *SD* = 25.11) compared to controls (*M* = 76.93, *SD* = 47.76), *t*(15) = 3.18, P < 0.006. To further examine whether functional changes in this cluster were associated with behavioural performance, we again conducted a series of Pearson’s correlations. Upon removal of an outlier (see “Additional Scene Cluster Results” in Supplementary for further detail), there was a moderate postive relationship between the COPE values extracted from this cluster and the dprime scores for scene stimuli, *r*(27) = 0.482, *P* = 0.008. Further examation revealed that a significant correlation was only apparent for the TBI group, *r*(15) = 0.574, *P* = 0.016, and not healthy controls, *r*(10) = 0.318, *P* = 0.316.

## Discussion

The present study focused on determining the role of temporal lobe activity in episodic memory behaviour following TBI. We showed for the first time, using converging evidence from behavioural and fMRI data, that episodic memory impairment following TBI appeared to be category-specific and related specific sub-regions within the temporal lobes. Deficits were most apparent for faces; TBI patients displayed reduced transverse temporal gyrus activation during face encoding and subsequent impairment on face recognition. TBI patients also displayed reduced fusiform gyrus activation for scenes; this is despite no statistically significant difference between groups during scene recognition. Interestingly, brain activation during face and scene encoding correlated with subsequent recognition for TBI patients but not in healthy control participants. Overall, these findings suggest that TBI: a) preferentially impairs episodic memory in specific domains, and b) aberrant neural processing may not be reflected in statistically significant differences behavioural assessments, and thus neural activation conveys complementary information undetected through examination of overt behaviour.

Broadly speaking, our findings are similar to those of previous studies of amnestic patients who show stimulus-sensitive impairments for complex stimuli such as faces and scenes (Taylor *et al.* 2007; Mundy *et al.* 2013). Our findings are also in concordance with a previous study demonstrating impaired face recognition in the TBI population (Valentine *et al.* 2006). Valentine *et al.* (2006) subjected participants to a range of facial recognition and learning tasks and found that while performance varied, deficits were more apparent for tasks with greater demands. More specifically, the most sensitive tasks were those which contained a larger number of faces to be encoded or had fewer presentations of the stimuli. Our task was comparably difficult in that participants were presented with a similar number of face stimuli which were only shown twice during the encoding phase; thus, it was not surprising we obtained a similar finding.

As expected, we found that the TBI group was generally slower than healthy controls in their reaction times. Both groups, however, responded more quickly to faces than to animals and scenes. This result somewhat aligns with a study conducted by Keightley et al. (2011) who found that participants reacted quicker to faces than scenes, despite accuracy being better for scene stimuli. We further explored whether speed-accuracy trade off could account for our findings, given participants displayed the poorest accuracy for face stimuli. We found that including reaction time as a covariate when determining between-group differences in face accuracy did not change the result. Instead, the rapid response to faces suggests that individuals with TBI may have performed superficial encoding of face stimuli, thus negatively impacting decision-making during recognition.

A unique aspect of our study was the inclusion of animal and scene stimuli, in addition to faces, which allowed us to investigate the effects of stimulus complexity on episodic memory. Interestingly, we found the TBI group displayed altered neural activity without showing statistically significant impairment of behavioural performance for scene stimuli, whereas changes on both functional and behavioural measures were apparent for faces. One potential reason for these results is that faces are more visually complex than scenes. Indeed, facial processing is a complex phenomenon requiring multifaceted processes across widespread cortical areas (Haxby et al. 2000). Unlike most other visual stimuli they are processed in a holistic and configural manner (Maurer et al. 2002; Park et al. 2009); thus, discrimination requires attention to detail and subtle perception of variable facial features (Haxby *et al.* 2000).

Our fMRI results provide further insights into the mechanisms that may underlie behavioural deficits for faces. We found that the TBI group showed a reduced response in the right transverse temporal gyrus extending to the planum temporale during encoding of faces. However, these areas do not form part of the core or extended face network (Haxby *et al.* 2002). The transverse temporal gyrus is predominantly implicated in auditory processing (Kaas et al. 1999; Warrier et al. 2009), although studies have also demonstrated its role in spontaneous inner speech (Hurlburt et al. 2016). The planum temporale has been shown to be involved in language functions (Shapleske et al. 1999). Therefore, it is possible that difference in BOLD activity in this temporal lobe region may reflect verbal strategy use. Interestingly, activation of this region correlated with behavioural performance only for individuals with TBI. It may be that individuals with TBI may therefore have made less use of verbal strategies when processing faces. This further supports the hypothesis that TBI participants encoded these stimuli in a rapid and superficial manner. In contrast, healthy controls did not display such an association, suggesting other strategies were used to aid encoding of faces.

Our other key finding was that the TBI group demonstrated reduced brain activity during encoding of scenes in the right posterior fusiform gyrus. The right posterior fusiform gyrus responds non-selectively to faces and scenes and generally may be involved with processing complex visual stimuli (Nakamura et al. 2000). This result suggests impaired recruitment of this temporal lobe sub-region during processing of scenes. Indeed, we found that individuals with TBI who had higher activation within this region performed better during scene recognition. Despite differences in brain activation, the groups did not differ with respect to behaviour. This finding may be due to scene stimuli containing a greater number of contextual cues that could further aid encoding and recollection. For example, individuals may have used cues such as the location (e.g. kitchen) or remembered certain salient scene features (e.g. item/s contained in the scene).

Although our findings show a clear brain-behaviour relationship, these associations oppose our initial hypotheses, which predicted that individuals with TBI would display greater temporal lobe activity in support of previous studies (e.g. Russell *et al.* 2011; Arenth *et al.* 2012; Gillis and Hampstead 2015). One general model that could explain this finding is that of cortical reorganisation following injury (Christodoulou et al. 2001; Levine et al. 2002; see Hillary 2008 for a review). That is, the pattern of cortical activation reflecting neural compensation or recovery following TBI is likely to depend on the length of time since an individual’s injury (Munoz-Cespedes et al. 2005). A key methodological difference is that our TBI participants were recruited at an average of 2 months’ post-injury whereas those in past studies were recruited over 1 year post-injury (Russell *et al.* 2011; Arenth *et al.* 2012; Gillis and Hampstead 2015). In a key study, Sanchez-Carrion et al. (2008) characterised the longitudinal changes in brain activity following TBI. They did so by assessing brain activity during a working memory task at 6 months and 1 year following the injury. Their findings show an initial reduction in brain activity at the 6 month time-points which gradually resolves by 1 year following the injury. Thus, discrepancies in our findings from past studies may reflect differences in recovery phases.

Our study has several important implications. From a clinical perspective, it is generally acknowledged that individuals have generalised episodic memory deficits after injury. Our findings provide evidence to the contrary and show that impairment is more apparent with complex visual stimuli such as faces and scenes. An obvious clinical translation is the need to provide strategies that promote deeper processing to better aid memory for these and other complex stimuli. In addition, our study provides further support for the utility of fMRI as a complementary source of information by demonstrating evidence of aberrant neural processing that was less evident in behavioural performance. This is important considering that most assessments of memory are based on behavioural performance.

There were some limitations in our study, however. Our episodic memory paradigm only allowed us to investigate functional activity during stimulus encoding. Therefore, we could not comment on the extent in which temporal structures are implicated in recognition of episodic stimuli following TBI. This may be an avenue for exploration in future research. Although not a limitation per se, we took an a-priori approach specifically focusing on the temporal lobes. While this allowed us to answer specific questions about the temporal lobes’ role in processing of episodic stimuli, it also limited our investigation of non-temporal contributions to episodic memory. For example, there is value in investigating the interaction between frontal and temporal regions to support encoding of more complex stimuli, especially considering the frontal lobes’ role in strategy, allocation of resources, and planning (Stuss and Alexander 2005; Vakil et al. 2019).

In conclusion, we found evidence demonstrating that individuals with TBI show impairment of episodic memory for complex stimuli and that this was associated with functional changes.

In comparison to healthy controls, we found that the TBI group displayed reduced activation in the right transverse temporal gyrus and fusiform gyrus during face and scene processing, respectively. We found that brain activation in these temporal lobe sub-regions were associated with behavioural performance for the TBI group and not healthy controls. These findings may be explained in terms of differences in strategy use and cortical reorganisation. Overall, we provide preliminary evidence demonstrating that following TBI: a) episodic memory impairment is domain-specific and more broadly dependent on the complexity of the stimuli, and b) aberrant neuronal activity may exist despite lack of evidence of significant impairment in behavioural performance, and therefore neural activation may be a more robust early indicator than behaviour.

## Abbreviations

TBI: traumatic brain injury
fMRI: functional magnetic resonance imaging
GSC: Glasgow Coma Scale
PTA: Post Traumatic Amnesia
WPTAS: Westmead Post Traumatic Amnesia Scale
DAI: diffuse axonal injury
EDH: extradural haematoma
ICH: intracerebral haemorrhage
SAH: subarachnoid haemorrhage
SDH: subdural haemorrhage
NAD: no abnormality detected
TR: repetition time
TE: echo time
COPE: contrast of parameter estimates

## Acknowledgements

This work was supported by the Multi-modal Australian ScienceS Imaging and Visualisation Environment (MASSIVE) HPC facility (www.massive.org.au). In addition, the authors would like to thank the staff at the Acquired Brain Injury Ward at Epworth Hospital (Richmond) and Bridge Road Imaging. The authors would also like to thank the participants who took part in the study.

## Supplementary Materials

**Supplementary Table 1.**
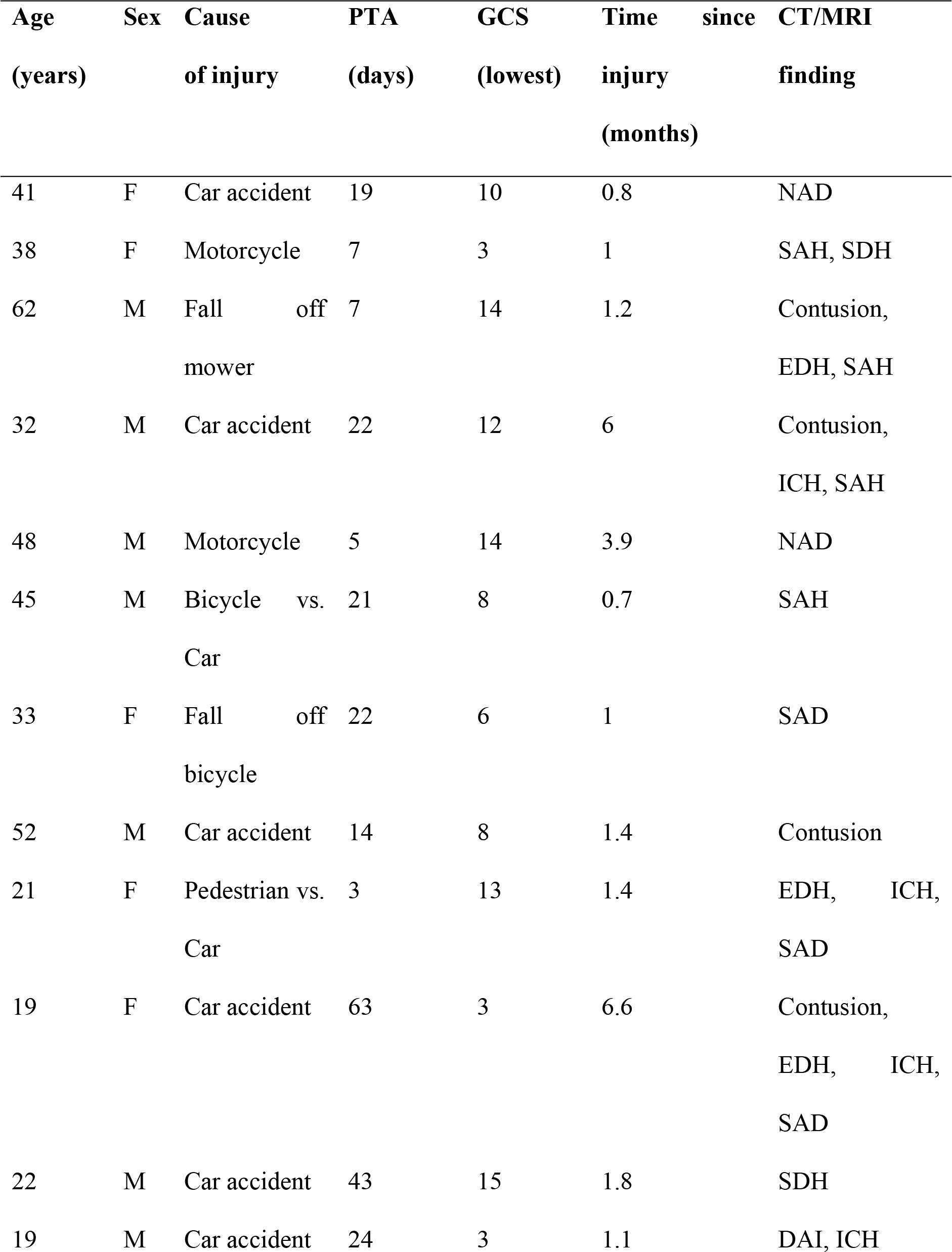
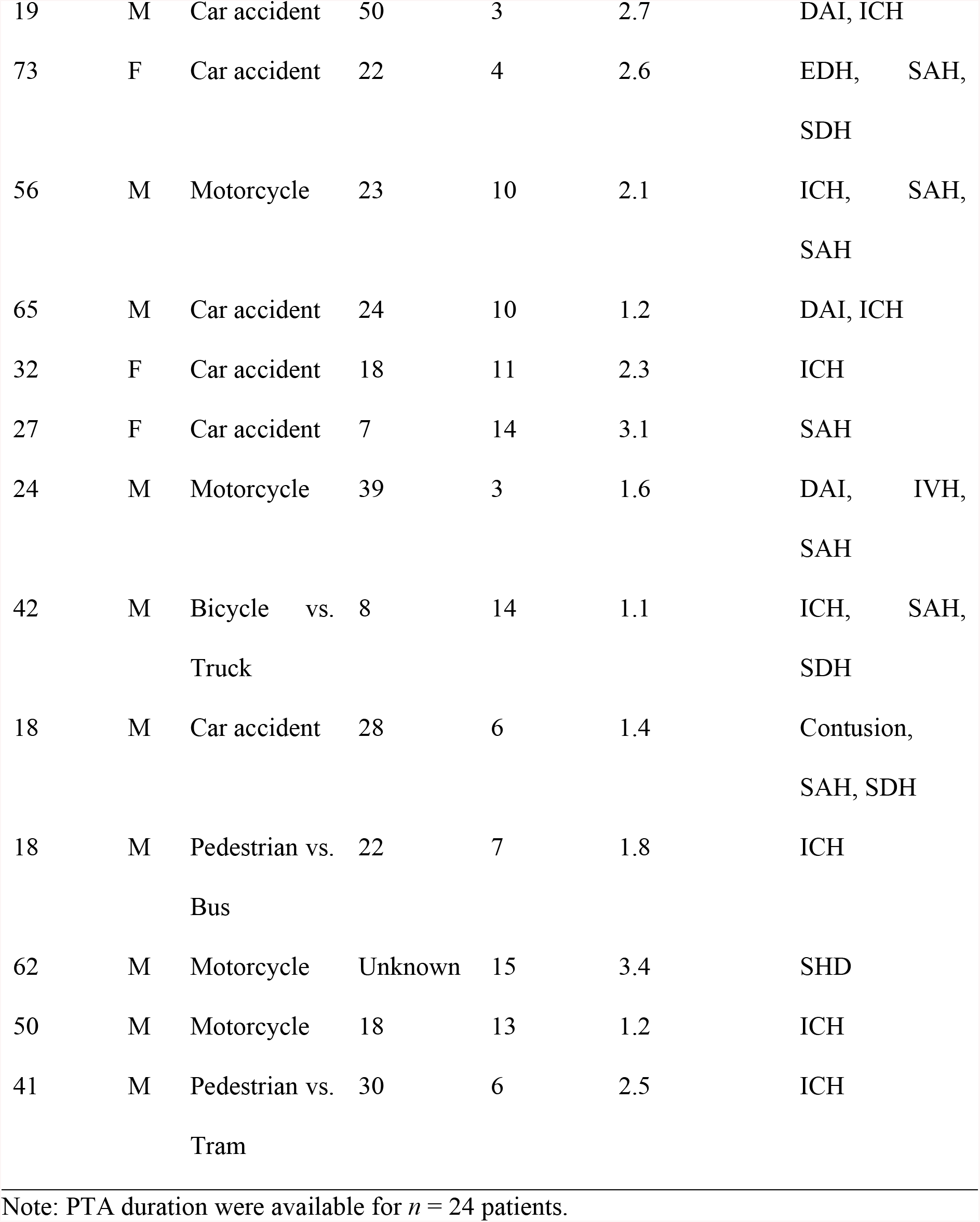
Clinical characteristics of each participant in the TBI group.

## Confidence Rating Analysis

To assess whether there were interactions between stimulus category, confidence, and correctness (i.e. correct/incorrect), linear mixed model was used. The outcome measure was the number of observations in each of the stimulus x confidence x correctness interactions. A model was fitted with stimulus category, confidence, correctness, group, and their interaction (stimulus category x confidence x correctness x group) as fixed effects, and participant as a random effect. Overall, there were more correct responses than incorrect, (95% CI, 0.27 – 12.27; *P* = 0.040). There was also a confidence x hit interaction (95% CI, 8.88 – 25.85; *P* < 0.001), such that there were more correct responses regardless of whether individuals were confident (95% CI, 31.53 – 40.02; *P* < 0.001) or not confident (95% CI, 14.94 – 23.42; *P* < 0.001). However, and more importantly, there were no significant 3-or 4-way interactions (p > 0.05); therefore, we felt justified in our decision to forego confidence ratings in our main analysis.

## Temporal Lobe Mask

**Supplementary Figure 1.**
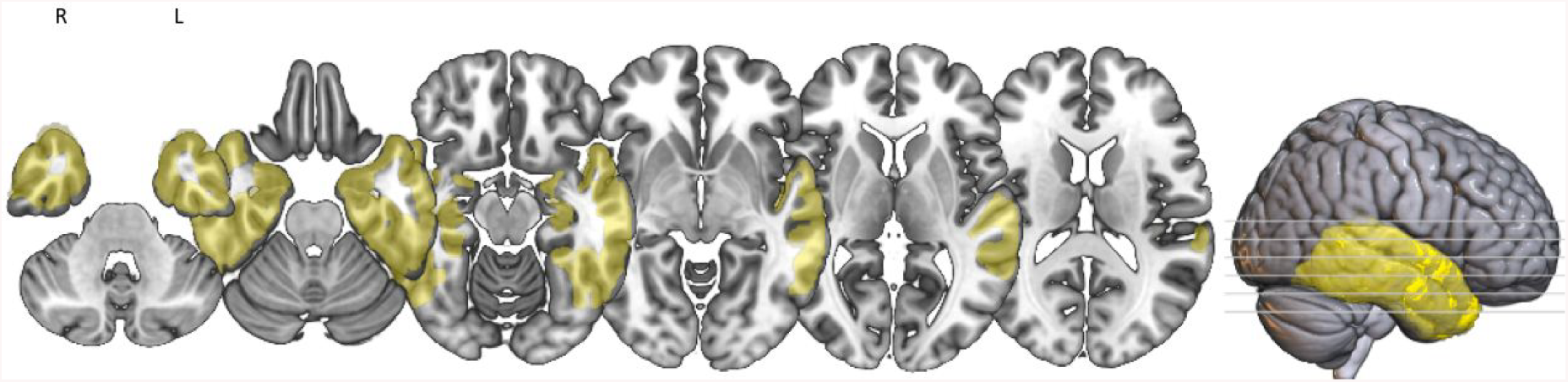
Temporal lobe mask (yellow) used in the group level analysis. The mask was generated from MNI Structural Atlas.

## Detailed MRI Preprocessing

Preprocessing of MRI data involved the following: each T1-weighted (T1w) image was corrected for intensity non-uniformity (INU) with N4BiasFieldCorrection (Tustison et al., 2010) and skull-stripped with a Nipype implementation of the antsBrainExtraction.sh workflow (using OASIS30ANTs as target template). Brain tissue segmentation of cerebrospinal fluid (CSF), white-matter (WM) and grey-matter (GM) was performed on the brain-extracted T1w using FAST (FSL 5.0.9; Zhang, Brady, & Smith, 2001). Cortical brain surfaces were reconstructed using recon-all (FreeSurfer 6.0.1; Dale, Fischl, & Sereno, 1999). Volume-based spatial normalisation to two standard spaces (MNI152NLin6Asym, MNI152NLin2009cAsym) was performed through nonlinear registration with antsRegistration (ANTs 2.2.0).

Functional data were skull-stripped using a custom methodology of fMRIPrep. fMRIPrep’s fieldmap-less approach was used to correct for susceptibility distortion using a deformation field resulting from co-registering the BOLD reference to the same-subject T1w-reference with its intensity inverted (Huntenburg, 2014; Wang et al., 2017).

The BOLD reference was then co-registered to the T1w reference using boundary-based registration (Greve & Fischl, 2009) with six degrees of freedom with bbregister (FreeSurfer). Head-motion parameters with respect to the BOLD reference (transformation matrices, and six corresponding rotation and translation parameters) were estimated using FSL’s MCFLIRT (Jenkinson, Bannister, Brady, & Smith, 2002). The BOLD time-series were resampled onto their original, native space by applying a single, composite transform to correct for head-motion and susceptibility distortions. The BOLD time-series were resampled into MNI152NLin6Asym standard space.

Non-steady state volumes were removed from preprocessed BOLD images and spatial smoothing with an isotropic, Gaussian kernel of 6mm FWHM (full-width half-maximum) was applied. Several confounding time-series were calculated based on the preprocessed BOLD: framewise displacement (FD), DVARS and three region-wise global signals. FD and DVARS are calculated, both using their implementations in Nipype (Power et al., 2014). The three global signals are extracted within the CSF, the WM, and the whole-brain masks. Additionally, a set of physiological regressors were extracted to allow for component-based noise correction (CompCor; Behzadi, Restom, Liau, & Liu, 2007). Principal components are estimated after high-pass filtering the preprocessed BOLD time-series (using a discrete cosine filter with 128s cut-off) for the two CompCor variants: temporal (tCompCor) and anatomical (aCompCor). tCompCor components are then calculated from the top 5% variable voxels within a mask covering the subcortical regions. For aCompCor, components are calculated within the intersection of the aforementioned mask and the union of CSF and WM masks calculated in T1w space, after their projection to the native space of each functional run (using the inverse BOLD-to-T1w transformation). Components are also calculated separately within the WM and CSF masks.

Many internal operations of fMRIPrep use Nilearn 0.6.1 (Abraham et al., 2014), mostly within the functional processing workflow. For more details of the pipeline, see https://fmriprep.readthedocs.io/en/1.0.8/workflows.html.

**Supplementary Table 2.** Regions of significant activation during encoding of faces.

| Contrast | Region | Voxels | Peak coordinates |  |  | *p*-value |
| --- | --- | --- | --- | --- | --- | --- |
|  |  |  | X | Y | Z |  |
| All | Right inferior occipital gyrus | 1030 | 44 | -68 | -14 | < 0.001 |
|  | Left fusiform gyrus | 626 | -38 | -40 | -26 | < 0.001 |
|  | Left fusiform gyrus | 48 | -34 | -10 | -28 | 0.015 |
|  | Right fusiform gyrus | 40 | 32 | -4 | -34 | 0.028 |
|  | Right hippocampus | 35 | 18 | -6 | -16 | 0.041 |
| HC > TBI | Right transverse temporal gyrus | 40 | 58 | -14 | 6 | 0.026 |
| TBI | Right fusiform gyrus | 810 | 40 | -50 | -22 | < 0.001 |
|  | Left fusiform gyrus | 475 | -38 | -42 | -26 | < 0.001 |
|  | Left fusiform gyrus | 33 | -36 | -12 | -32 | 0.030 |
| HC | Right fusiform gyrus | 305 | 46 | -44 | -16 | < 0.001 |
|  | Right fusiform gyrus | 86 | 38 | -6 | -34 | 0.002 |
|  | Right fusiform gyrus | 48 | 54 | -18 | 10 | 0.021 |

**Supplementary Table 3.**
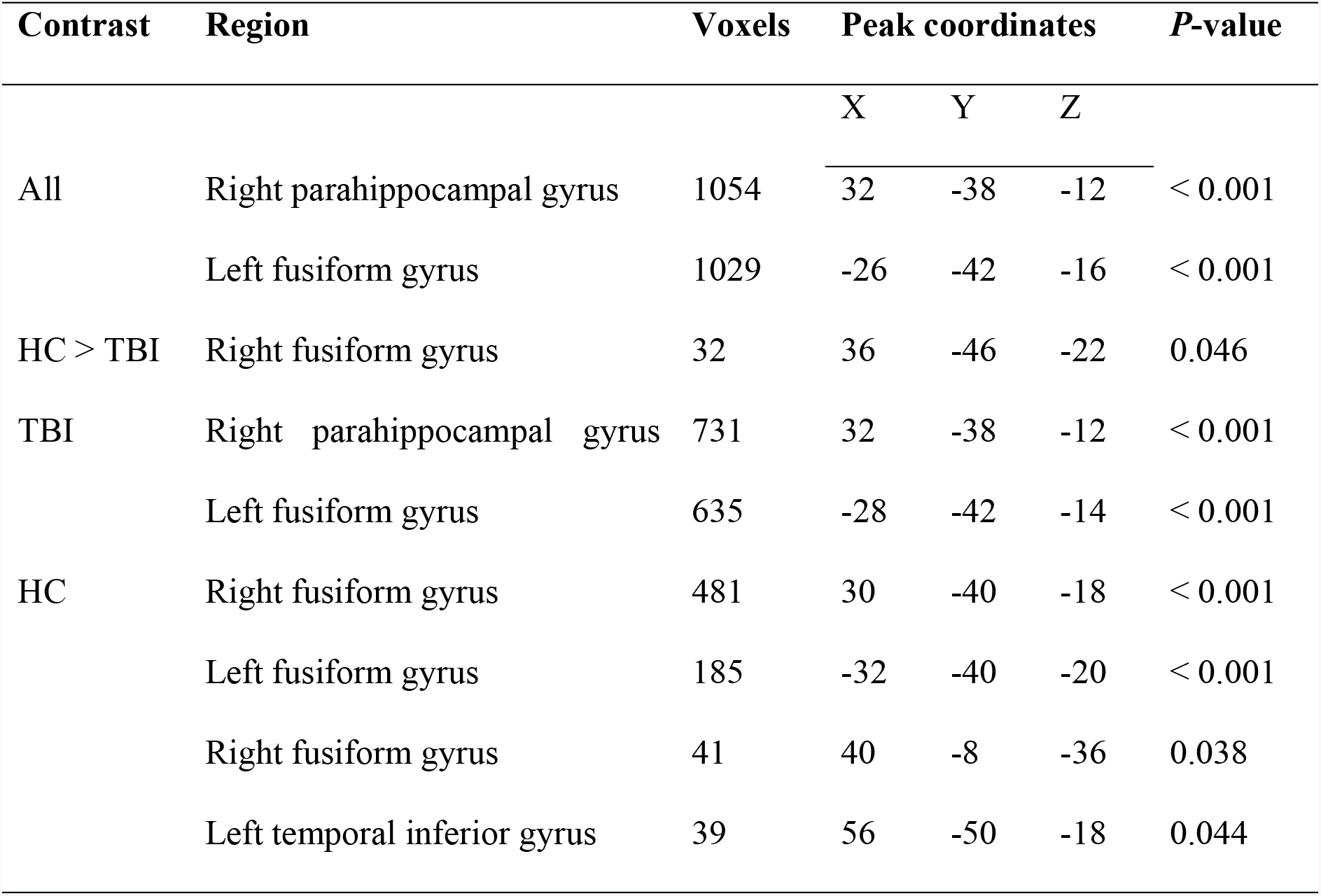
Regions of significant activation during encoding of scenes.

**Supplementary Table 4.** Regions of significant activation during encoding of animals.

| Contrast | Region | Voxels | Peak coordinates |  |  | P-value |
| --- | --- | --- | --- | --- | --- | --- |
|  |  |  | X | Y | Z |  |
| All | Right fusiform gyrus | 1511 | 44 | -60 | -18 | < 0.001 |
|  | Left inferior occipital gyrus | 1294 | -44 | -72 | -16 | < 0.001 |
|  | Right fusiform gyrus | 39 | 32 | -8 | -32 | 0.031 |
| TBI | Right fusiform gyrus | 1178 | 44 | -58 | -22 | < 0.001 |
|  | Left fusiform gyrus | 995 | -46 | -60 | -14 | < 0.001 |
| HC | Right fusiform gyrus | 676 | 48 | -62 | -12 | < 0.001 |
|  | Left fusiform gyrus | 176 | -48 | -58 | -10 | < 0.001 |
|  | Left fusiform gyrus | 82 | -50 | -46 | -24 | 0.004 |
|  | Right temporal inferior gyrus | 78 | 44 | 0 | -40 | 0.005 |
|  | Left fusiform gyrus | 52 | -38 | -70 | -16 | 0.024 |

## Additional Scene Cluster Results

The results reported in the main text are after removal of an outlier, defined as a data point 3.29 standard deviations from the mean (Tabachnick, Fidell, & Ullman, 2007). Retaining this outlier would have still resulted in a significant correlation between COPE value and the dprime scores for scenes overall, *r*(27) = 0.436, *P* = 0.016. However, the correlation would have only trended towards significance for the TBI group *r*(15) = 0.453, *P* = 0.059.

**Supplementary Figure 2.**
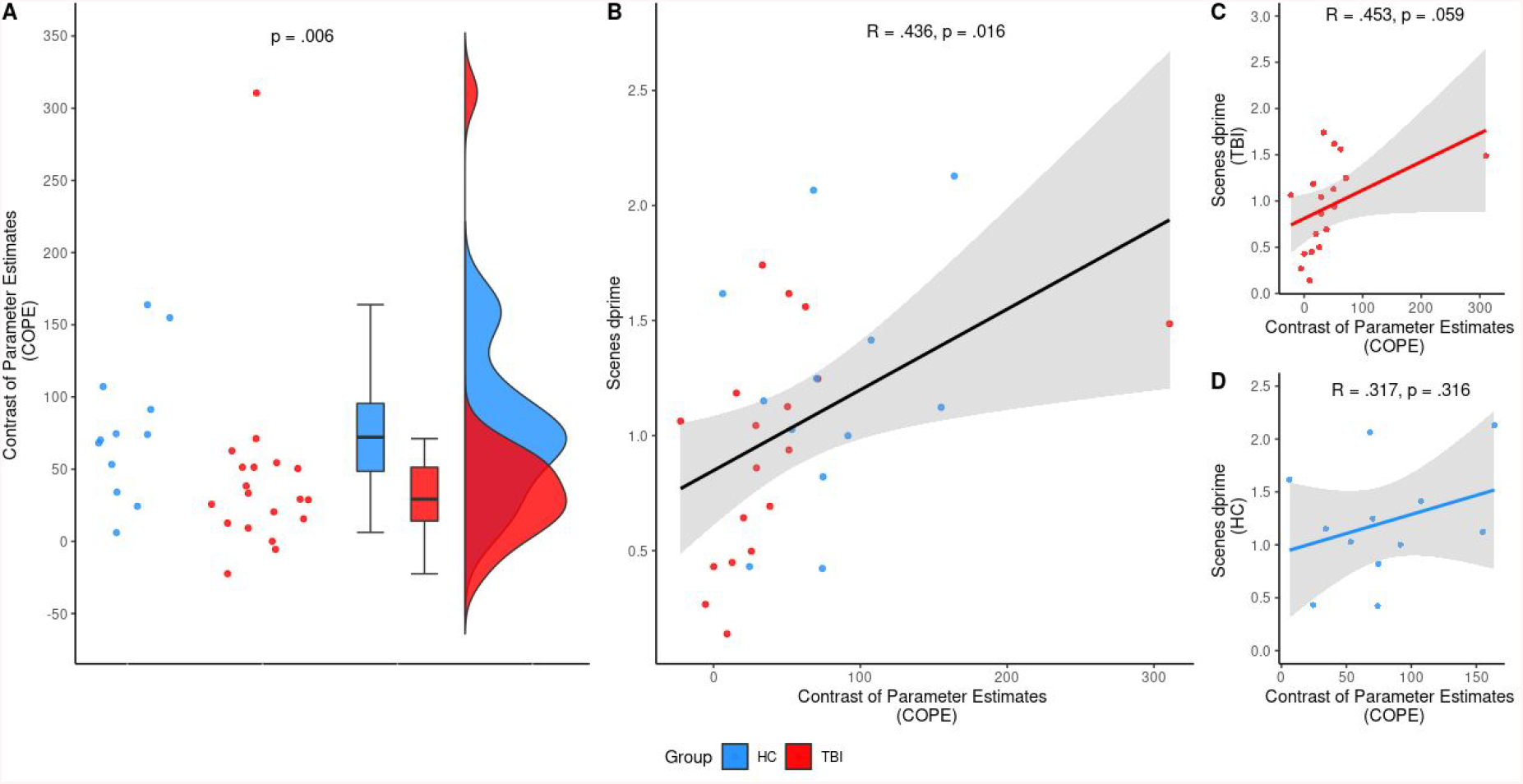
Results for the scene cluster if the outlier was retained. A) Plot of the COPE extracted from the significant cluster in the right posterior fusiform gyrus. The outlier would have affected the results such that there was no difference in the COPE between the TBI group (red) and healthy controls (blue; *P* = 0.128). B) Overall, there was a signification association between the COPE and the dprime for scene stimuli (*P* = 0.018). C) There was a trend towards a significant correlation for the TBI group between the COPE and the dprime for scene stimuli (*P* = 0.059). D) There was no significant correlation for the healthy controls (*P* = 0.316).

## References

Arenth PM, Russell KC, Scanlon JM, Kessler LJ, Ricker JH. 2012. Encoding and recognition after traumatic brain injury: neuropsychological and functional magnetic resonance imaging findings. J Clin Exp Neuropsychol. 34:333–344.

Ariza M, Serra-Grabulosa JM, Junque C, Ramirez B, Mataro M, Poca A, Bargallo N, Sahuquillo J. 2006. Hippocampal head atrophy after traumatic brain injury. Neuropsychologia. 44:1956–1961.

Azouvi P, Arnould A, Dromer E, Vallat-Azouvi C. 2017. Neuropsychology of traumatic brain injury: An expert overview. Rev Neurol (Paris). 173:461–472.

Barlow KM. 2013. Traumatic brain injury. Handb Clin Neurol. 112:891–904.

Bigler ED. 2001. The lesion (s) in traumatic brain injury: Implications for clinical neuropsychology. Archives of clinical neuropsychology. 16:95–131.

Bigler ED, Johnson SC, Anderson CV, Blatter DD, Gale SD, Russo AA, Ryser DK, Macnamara SE, Bailey BJ, Hopkins RO. 1996. Traumatic brain injury and memory: The role of hippocampal atrophy. Neuropsychology. 10:333.

Cameron KA, Yashar S, Wilson CL, Fried I. 2001. Human hippocampal neurons predict how well word pairs will be remembered. Neuron. 30:289–298.

Christodoulou C, DeLuca J, Ricker J, Madigan N, Bly B, Lange G, Kalnin A, Liu W, Steffener J, Diamond B. 2001. Functional magnetic resonance imaging of working memory impairment after traumatic brain injury. Journal of Neurology, Neurosurgery & Psychiatry. 71:161–168.

Daneshvar DH, McKee AC. 2015. Traumatic Brain Injury. 219–235.

Dickerson BC, Eichenbaum H. 2010. The episodic memory system: neurocircuitry and disorders. Neuropsychopharmacology. 35:86–104.

Downing PE, Chan AW, Peelen MV, Dodds CM, Kanwisher N. 2006. Domain specificity in visual cortex. Cereb Cortex. 16:1453–1461.

Draper K, Ponsford J. 2008. Cognitive functioning ten years following traumatic brain injury and rehabilitation. Neuropsychology. 22:618–625.

Eichenbaum H. 2017. Prefrontal-hippocampal interactions in episodic memory. Nat Rev Neurosci. 18:547–558.

Epstein RA, Ward EJ. 2010. How reliable are visual context effects in the parahippocampal place area? Cereb Cortex. 20:294–303.

Gillis MM, Hampstead BM. 2015. A two-part preliminary investigation of encoding-related activation changes after moderate to severe traumatic brain injury: hyperactivation, repetition suppression, and the role of the prefrontal cortex. Brain Imaging Behav. 9:801–820.

Graham KS, Barense MD, Lee AC. 2010. Going beyond LTM in the MTL: a synthesis of neuropsychological and neuroimaging findings on the role of the medial temporal lobe in memory and perception. Neuropsychologia. 48:831–853.

Haxby JV, Hoffman EA, Gobbini MI. 2000. The distributed human neural system for face perception. Trends in cognitive sciences. 4:223–233.

Haxby JV, Hoffman EA, Gobbini MI. 2002. Human neural systems for face recognition and social communication. Biological psychiatry. 51:59–67.

Hillary FG. 2008. Neuroimaging of working memory dysfunction and the dilemma with brain reorganization hypotheses. J Int Neuropsychol Soc. 14:526–534.

Hoffman EA, Haxby JV. 2000. Distinct representations of eye gaze and identity in the distributed human neural system for face perception. Nature neuroscience. 3:80–84.

Hurlburt RT, Alderson-Day B, Kühn S, Fernyhough C. 2016. Exploring the ecological validity of thinking on demand: neural correlates of elicited vs. spontaneously occurring inner speech. PloS one. 11:e0147932.

Kaas JH, Hackett TA, Tramo MJ. 1999. Auditory processing in primate cerebral cortex. Current opinion in neurobiology. 9:164–170.

Keightley ML, Chiew KS, Anderson JA, Grady CL. 2011. Neural correlates of recognition memory for emotional faces and scenes. Soc Cogn Affect Neurosci. 6:24–37.

Kesler ML, Andersen AH, Smith CD, Avison MJ, Davis CE, Kryscio RJ, Blonder LX. 2001. Neural substrates of facial emotion processing using fMRI. Cognitive Brain Research. 11:213–226.

Levine B, Cabeza R, McIntosh A, Black S, Grady C, Stuss D. 2002. Functional reorganisation of memory after traumatic brain injury: a study with H2150 positron emission tomography. Journal of Neurology, Neurosurgery & Psychiatry. 73:173–181.

Maurer D, Le Grand R, Mondloch CJ. 2002. The many faces of configural processing. Trends in cognitive sciences. 6:255–260.

Moscovitch M, Cabeza R, Winocur G, Nadel L. 2016. Episodic Memory and Beyond: The Hippocampus and Neocortex in Transformation. Annu Rev Psychol. 67:105–134.

Mundy ME, Downing PE, Dwyer DM, Honey RC, Graham KS. 2013. A critical role for the hippocampus and perirhinal cortex in perceptual learning of scenes and faces: complementary findings from amnesia and fMRI. Journal of Neuroscience. 33:10490–10502.

Mundy ME, Honey RC, Downing PE, Wise RG, Graham KS, Dwyer DM. 2009. Material-independent and material-specific activation in functional MRI after perceptual learning. Neuroreport. 20:1397–1401.

Munoz-Cespedes JM, Rios-Lago M, Paul N, Maestu F. 2005. Functional neuroimaging studies of cognitive recovery after acquired brain damage in adults. Neuropsychol Rev. 15:169–183.

Muschelli J, Nebel MB, Caffo BS, Barber AD, Pekar JJ, Mostofsky SH. 2014. Reduction of motion-related artifacts in resting state fMRI using aCompCor. Neuroimage. 96:22–35.

Nakamura K, Kawashima R, Sato N, Nakamura A, Sugiura M, Kato T, Hatano K, Ito K, Fukuda H, Schormann T. 2000. Functional delineation of the human occipito-temporal areas related to face and scene processing: a PET study. Brain. 123:1903–1912.

Nakase-Richardson R, Sherer M, Seel RT, Hart T, Hanks R, Arango-Lasprilla JC, Yablon SA, Sander AM, Barnett SD, Walker WC, Hammond F. 2011. Utility of post-traumatic amnesia in predicting 1-year productivity following traumatic brain injury: comparison of the Russell and Mississippi PTA classification intervals. J Neurol Neurosurg Psychiatry. 82:494–499.

Park J, Newman LI, Polk TA. 2009. Face processing: the interplay of nature and nurture. The Neuroscientist. 15:445–449.

Rabinowitz AR, Levin HS. 2014. Cognitive sequelae of traumatic brain injury. Psychiatr Clin North Am. 37:1–11.

Rogers TT, Hocking J, Mechelli A, Patterson K, Price C. 2005. Fusiform activation to animals is driven by the process, not the stimulus. Journal of Cognitive Neuroscience. 17:434–445.

Russell KC, Arenth PM, Scanlon JM, Kessler LJ, Ricker JH. 2011. A functional magnetic resonance imaging investigation of episodic memory after traumatic brain injury. J Clin Exp Neuropsychol. 33:538–547.

Sanchez-Carrion R, Fernandez-Espejo D, Junque C, Falcon C, Bargallo N, Roig T, Bernabeu M, Tormos JM, Vendrell P. 2008. A longitudinal fMRI study of working memory in severe TBI patients with diffuse axonal injury. Neuroimage. 43:421–429.

Shapleske J, Rossell SL, Woodruff P, David A. 1999. The planum temporale: a systematic, quantitative review of its structural, functional and clinical significance. Brain Research Reviews. 29:26–49.

Shores EA, Marosszeky J, Sandanam J, Batchelor J. 1986. Preliminary validation of a clinical scale for measuring the duration of post-traumatic amnesia. Med J Aust. 144:569–572.

Simons JS, Spiers HJ. 2003. Prefrontal and medial temporal lobe interactions in long-term memory. Nat Rev Neurosci. 4:637–648.

Stuss DT, Alexander MP. 2005. Does damage to the frontal lobes produce impairment in memory? Current Directions in Psychological Science. 14:84–88.

Taylor KJ, Henson RN, Graham KS. 2007. Recognition memory for faces and scenes in amnesia: dissociable roles of medial temporal lobe structures. Neuropsychologia. 45:2428–2438.

Tittle A, Burgess GH. 2011. Relative contribution of attention and memory toward disorientation or post-traumatic amnesia in an acute brain injury sample. Brain Injury. 25:933–942.

Tulving E. 2002. Episodic memory: from mind to brain. Annual review of psychology. 53:1–25.

Vakil E. 2005. The effect of moderate to severe traumatic brain injury (TBI) on different aspects of memory: a selective review. J Clin Exp Neuropsychol. 27:977–1021.

Vakil E, Greenstein Y, Weiss I, Shtein S. 2019. The Effects of Moderate-to-Severe Traumatic Brain Injury on Episodic Memory: a Meta-Analysis. Neuropsychol Rev. 29:270–287.

Valentine T, Powell J, Davidoff J, Letson S, Greenwood R. 2006. Prevalence and correlates of face recognition impairments after acquired brain injury. Neuropsychol Rehabil. 16:272–297.

Wagner AD, Shannon BJ, Kahn I, Buckner RL. 2005. Parietal lobe contributions to episodic memory retrieval. Trends Cogn Sci. 9:445–453.

Warrier C, Wong P, Penhune V, Zatorre R, Parrish T, Abrams D, Kraus N. 2009. Relating structure to function: Heschl’s gyrus and acoustic processing. Journal of Neuroscience. 29:61–69.

